# A longevity-specific bank of induced pluripotent stem cells from centenarians and their offspring

**DOI:** 10.1101/2024.03.12.584663

**Authors:** Todd W. Dowrey, Samuel F. Cranston, Nicholas Skvir, Yvonne Lok, Brian Gould, Bradley Petrowitz, Daniel Villar, Jidong Shan, Marianne James, Mark Dodge, Anna C. Belkina, Richard M. Giadone, Paola Sebastiani, Thomas T. Perls, Stacy L. Andersen, George J. Murphy

## Abstract

Centenarians provide a unique lens through which to study longevity, healthy aging, and resiliency. Moreover, models of *human* aging and resilience to disease that allow for the testing of potential interventions are virtually non-existent. We obtained and characterized over 50 centenarian and offspring peripheral blood samples including those connected to functional independence data highlighting resistance to disability and cognitive impairment. Targeted methylation arrays were used in molecular aging clocks to compare and contrast differences between biological and chronological age in these specialized subjects. Isolated peripheral blood mononuclear cells (PBMCs) were then successfully reprogrammed into high-quality induced pluripotent stem cell (iPSC) lines which were functionally characterized for pluripotency, genomic stability, and the ability to undergo directed differentiation. The result of this work is a one-of-a-kind resource for studies of human longevity and resilience that can fuel the discovery and validation of novel therapeutics for aging-related disease.

## INTRODUCTION

Individuals with exceptional longevity (EL) age more successfully than the general population by extending their healthspan and decreasing the proportion of their lives spent with aging-related disease, deemed the ‘compression of morbidity’^1–4^. Centenarians (>100 years of age) provide a unique lens through which to study EL, healthy aging, and disease resilience and resistance^5,6^. In the last decade, several studies have provided evidence that centenarians exhibit delayed onset or escape aging-related diseases such as cancer, cardiovascular disease, and Alzheimer’s disease (AD) while markedly delaying disability^4,6,7^. Although recent work has identified genetic variants associated with healthful aging, insights into how these elements promote longevity remain unclear^8^. Understanding the regulatory networks that promote resistance to aging-related disease may provide mechanistic insights into this process and inform the development of therapeutics to slow or reverse aging. Problematically, however, models of *human* aging, longevity, and resistance to and/or resilience against disease that allow for the functional testing of potential interventions are virtually non-existent.

Induced pluripotent stem cells (iPSCs) faithfully capture the genetic background of the person from whom they are created and are revolutionizing pre-clinical drug screening by exhibiting the power of precision medicine. Beginning soon after their initial discovery, iPSCs have been used to model diseases as well as screen drugs for the treatment of amyotrophic lateral sclerosis^9^, spinal muscular atrophy^10^, and other neurodegenerative^11^ and muscular^12^ disorders. Notably and importantly, these model systems have been applied to and faithfully recapitulate disease pathologies associated with aging that manifest late in life, such as in AD^13–17^ and in our own work on familial ATTR-amyloidosis^18–21^. The versatility in these systems to produce any cell and tissue type of the body allows for interrogation of multiple, aging-impacted tissues, many of which have been demonstrated to age at different rates^22^. This potential utility of iPSCs in aging research has recently been highlighted as a platform for both longevity studies as well as drug discovery^23^. EL iPSCs also provide a complement for studies performed on aging disease-specific lines, such as those derived from patients with Progeria^24,25^, given their potential in understanding mechanisms that embolden resiliency and resistance to accelerated aging-related disease.

Here, we report a novel bank of longevity-specific peripheral blood mononuclear cells (PBMCs) and resultant iPSCs from subjects with EL. The result of this work is a highly characterized, one-of-a-kind resource that can aid in a host of aging-related studies. This flexible iPSC library represents a unique, permanent resource that can be harnessed by any investigator for molecular and functional analyses of resiliency, longevity, and aging. It also provides a much-needed human platform for the discovery of novel geroprotective agents and/or the validation of findings from other data banks, tissue repositories, or models. Lastly, our ability to connect detailed phenotypic data obtained from the subjects as well as molecular and biological aging data to the created lines enables the identification of those at the extremes of both physical and cognitive functionality for study.

## RESULTS

### Identification and characterization of individuals displaying exceptional longevity

PBMCs were collected from 36 centenarians (mean age at last contact/death 104.58 ± 3.5 years, 56% females), 16 offspring (mean age at last contact 72.8 ± 7.3 years, range 60-86 years, 69% females), and 2 non-EL offspring spouses (85.5 ± 4.9 years, 0% females) (**Table 1**). Among the centenarians for whom cognitive and/or functional status could be determined at age 100, 73% (8/11) were cognitively healthy and 76% were Activities of Daily Living (ADL) independent at age 100 years. Among offspring with sufficient data to characterize current cognitive and/or functional status, 82% (9/11) were cognitively healthy and 100% (n=14) were independent in performing Instrumental Activities of Daily Living (IADLs) at the time of assessment. For investigators seeking additional protected phenotypic data on study participants, applications can be made to the ELITE portal data hub (https://eliteportal.synapse.org/) where this information is securely housed.

**Table 1.** Cognitive and functional status of EL subjects. Demographic (age, sex), cognitive, and functional status of subjects in this bank are organized by age bracket and separated by cohort (Centenarian, Offspring, and Offspring Spouses) with number of subjects listed next to each identifier. Cognitive status is determined by clinical consensus review of neuropsychological assessment scores or cognitive screeners detailed in the methods. Functional independence status was determined by performance on the Barthel Activities of Daily Living Index (ADLs) for centenarians and Instrumental Activities of Daily Living (IADLs) for offspring and spouses.

| Age at Draw | Sex (N) | Cognitively healthy at age 100 (N) | ADL independent at age 100 (N) |
| --- | --- | --- | --- |
| 100-104 | Male (10)<br>Female (11) | Yes (4)<br>No (3)<br>Unknown (14) | Yes (12)<br>No (2)<br>Unknown (7) |
| 105-109 | Male (6)<br>Female (7) | Yes (3)<br>No (0)<br>Unknown (10) | Yes (4)<br>No (3)<br>Unknown (6) |
| 110+ | Male (0)<br>Female (2) | Yes (1)<br>No (0)<br>Unknown (1) | Yes (1)<br>No (0)<br>Unknown (1) |

**Offspring**
| Age at Draw | Sex (N) | Cognitively healthy (N) | IADL independent (N) |
| --- | --- | --- | --- |
| 60-69 | Male (1)<br>Female (4) | Yes (3)<br>No (0)<br>Unknown (2) | Yes (5)<br>No (0)<br>Unknown (0) |
| 70-79 | Male (4)<br>Female (5) | Yes (6)<br>No (1)<br>Unknown (2) | Yes (9)<br>No (0)<br>Unknown (0) |
| 80+ | Male (0)<br>Female (2) | Yes (0)<br>No (1)<br>Unknown (1) | Yes (2)<br>No (0)<br>Unknown (0) |

**Table 1. Cognitive and functional status of EL subjects.**
| Age at Draw | Sex (N) | Cognitively healthy (N) | IADL independent (N) |
| --- | --- | --- | --- |
| 80+ | Male (2)<br>Female (0) | Yes (0)<br>No (0)<br>Unknown (2) | Yes (2)<br>No (0)<br>Unknown (0) |

### Comparisons of biological versus chronological age in EL subjects

Methylation profiling was performed with Illumina Infinium arrays on 18 subjects in this bank which allowed for estimation of biological age through established aging clocks, including the pan-tissue Horvath DNAmAge^26,27^ clock, Hannum^28^, DunedinPACE^29^, and Phenoage^30^ clocks, as well as a recently published aging clock trained on datasets that included a relatively high proportion of centenarians (ENCen40+)^31^ (**Figure 1**). This panel of epigenetic clocks comprises a wide array of models, trained on both chronological and phenotypic measurements, and spanning multiple generations of Illumina methylation chips. To facilitate compatibility between each of these models with the newest Illumina beadchips, imputation for the small subsets of missing CpGs for each model was performed where applicable (see methods). Despite differences in newer generation (Illumina EPICv2) versus older generation methylation arrays, estimates and model behavior seen with our samples were largely in line with expectations. Models trained on the older Illumina 27k/450k arrays saw accurate age prediction in younger individuals (Horvath median absolute error (MAE) = 3.67 / Pearson correlation coefficient (R) = 0.93, Hannum MAE = 4.42 / R = 0.94 on non-centenarian samples), as well as a general under-predictive trend in EL samples. Meanwhile, clocks measuring putative biological aging (PhenoAge, DunedinPACE) showed similarly consistent correlation. PhenoAge showed a correlation coefficient of 0.94 with chronological age, while DunedinPACE centered around an average value of 1 (representing the age acceleration of a healthy adult) with gradual increase in the age acceleration of chronologically older individuals (R=0.62) as noted by the authors in the original manuscript^29^. Based on the spectrum of clocks employed, the centenarians in this bank mapped biologically younger to varying extents. Notably, the ENCen40+ clock estimated an average 6.0 year reduction in biological age and a much higher average degree of accuracy in centenarians relative to other current models. These data reflected the relative health metrics and clinical history associated with each subject, with those displaying functional independence and higher cognitive function having the most pronounced reduction in biological age. Interestingly, EL offspring displayed more variable biological age status, with some subjects estimated to have a higher biological age as well as some estimated to have a lower biological age (**Figure 1**). The greater variation in the offspring may be the result of greater heterogeneity in the presence or absence of genetic signatures associated with EL. Of note, a subset of the PBMCs collected in this study were also characterized via single cell RNA sequencing and CITEseq analyses^32^, as well as a 40-color flow cytometry panel to extensively examine immune cell subtypes and identify unique features in EL subjects (**Supplementary Figures 1-2**).

**Figure 1.**
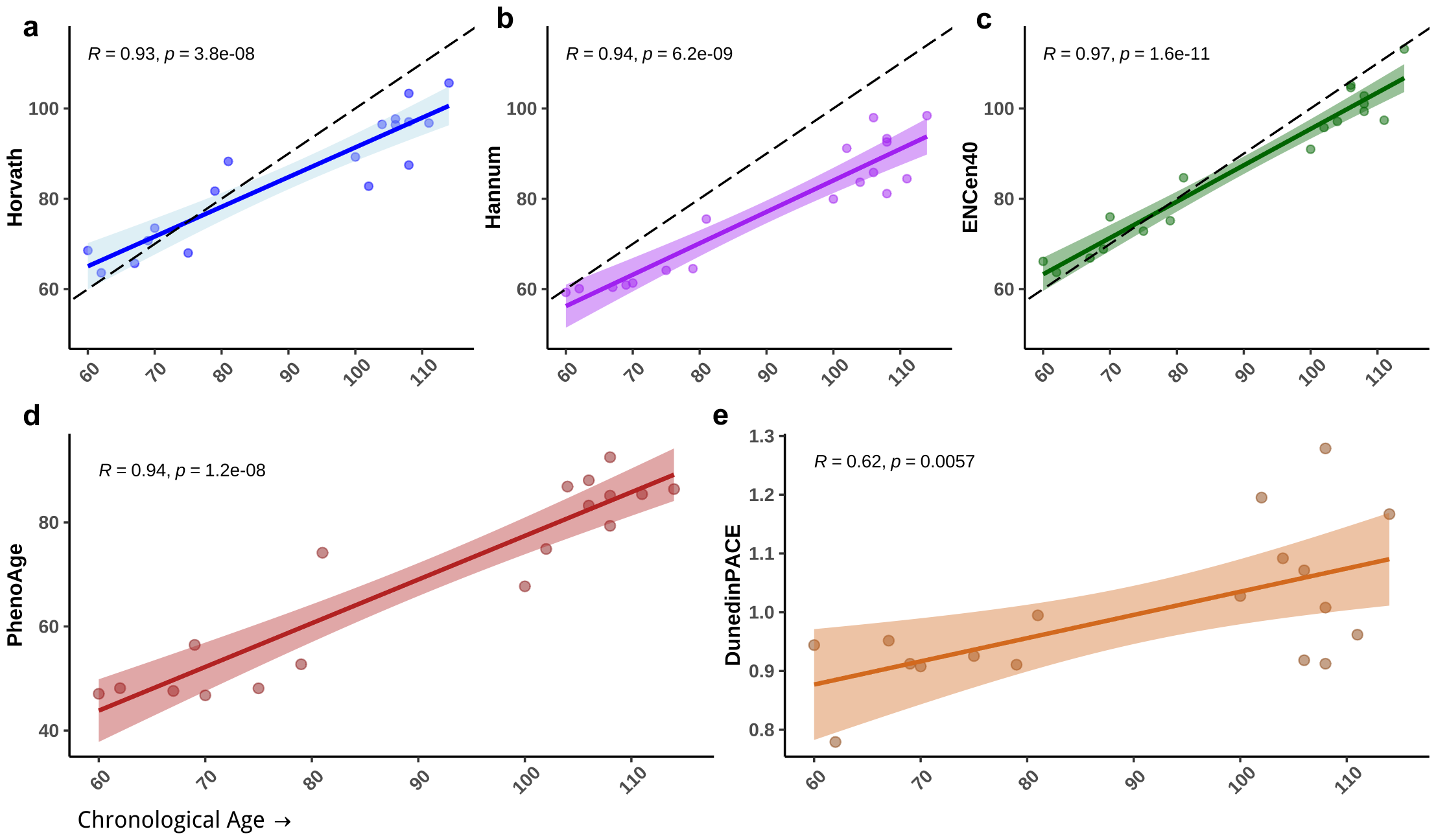
Comparisons of biological versus chronological age in EL subjects. Parallel age estimates from five well-established models of epigenetic aging for EL PBMCs. The colored lines represent the best linear fit from each model, while the dotted black lines show a theoretical perfect linear correlation for reference, where applicable. Each data point represents an individual within our bank. The top three panels show models trained on chronological age including the original 2013 pan-tissue Horvath (**a**), Hannum (**b**), and ENCen40+ (**c**) clocks. Panels **d** and **e** show predictions from models (PhenoAge and DunedinPACE) trained with the additional aid of clinical and phenotypic measurements. These clocks return more novel measurements that align more closely to a putative biological age, or rate of age acceleration, as opposed to chronological age.

### Establishment and characterization of a longevity-specific iPSC bank

Of the 54 subjects from which PBMCs were collected and isolated, 19 iPSC lines have been generated across EL sub-groups (**Table 2**). All lines were created using our established methodologies^33–36^ with at least three independent clones generated from each subject. All lines met stringent quality control parameters for pluripotency and functionality^33–36^ and were expanded to facilitate sharing in an opensource approach. Briefly, each line was characterized for pluripotency using TRA-1-81 expression (**Figure 2a**), as well as for genomic integrity via karyotype analysis (**Figure 2b**). Although otherwise karyotypically normal, 3 male centenarians displayed mosaic loss of y chromosome (**Figure 2c**). Lastly, teratoma assays were also performed to confirm pluripotency and the ability to generate tissue types representative of all three germ layers^34,37,38^ (**Figure 2d**). Importantly, regardless of subject age, no failures in iPSC generation or pluripotency competence were observed.

**Table 2.** Demographic information of EL and non-EL PBMC and iPSC lines. EL Subjects are classified by age group, sex, and EL status. Included in this table are centenarian offspring as well as non-EL controls in the offspring age group.

| **Age** | **Sex** | **PBMC**<br>**Collected** | **iPSC**<br>**Generated** |
| --- | --- | --- | --- |
| 100-104 | Male | 10 | 1 |
|  | Female | 11 | 2 |
| 105-109 | Male | 6 | 5 |
|  | Female | 7 | 5 |
| 110+ | Male | 0 | 0 |
|  | Female | 2 | 2 |
| EL<br>Offspring | Male | 5 | 1 |
|  | Female | 11 | 3 |
| Non-EL<br>controls | Male | 2 | 0 |
|  | Female | 0 | 0 |
| **Total** |  | **54** | **19** |

**Figure 2.**
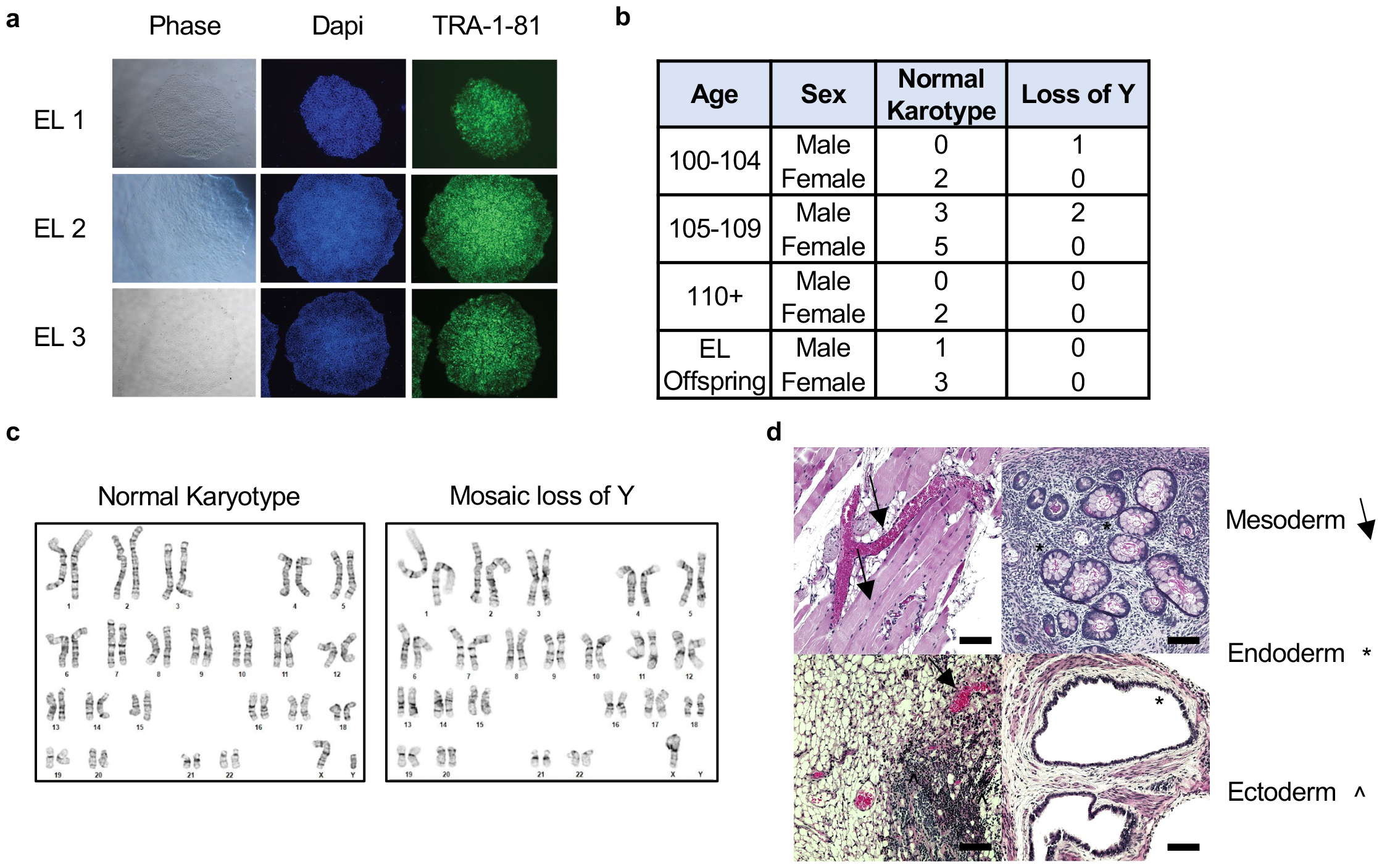
Representative characterization of EL-specific iPSCs. **a**) Representative images of iPSC lines under brightfield (left), DAPI (middle), and TRA-1-81 (right). Images taken at 10X magnification. **b**) Table of iPSC lines generated including demographic information (age, sex) and karyotype outcome based on g-band analyses. **c**) Representative karyotype reports of male EL iPSCs showing a normal karyotype (left) and mosaic loss of y chromosome (right). **d**) Representative images of teratoma mass hematoxylin and eosin stains from EL iPSC lines representing mesoderm (arrows), endoderm (asterisks) and ectoderm (accent) tissue. Scale bars: 100 µM.

### Forward programming of EL-specific iPSCs into cortical neurons

As a demonstration of the potential of the established EL-specific iPSC lines to undergo directed differentiation, and as neuronal cell types are impacted by many aging-related diseases with large socioeconomic burden, we conducted transcription factor-mediated differentiation to forebrain cortical neurons using established methods^39,40^. Briefly, cells were engineered to have doxycycline-inducible expression of the neuronal transcription factor neurogenin 2 (NGN2) via transposase-mediated integration. Upon a 3-day NGN2 induction using doxycycline, cultures comprised of neural progenitor cells that were replated onto a specialized matrix of laminin and fibronectin and allowed to mature for 14 days in neuronal BrainPhys (STEMCELL Technologies) media supplemented with growth factors NT3, BDNF, and B27. Resulting iPSC-derived neurons (iNeurons) displayed uniform morphology and expression of neuronal markers TUJ1^41,42^ and ISLET1/2^43,44^ (**Figure 3**). All lines tested performed comparably in their ability to efficiently generate iNeurons.

**Figure 3.**
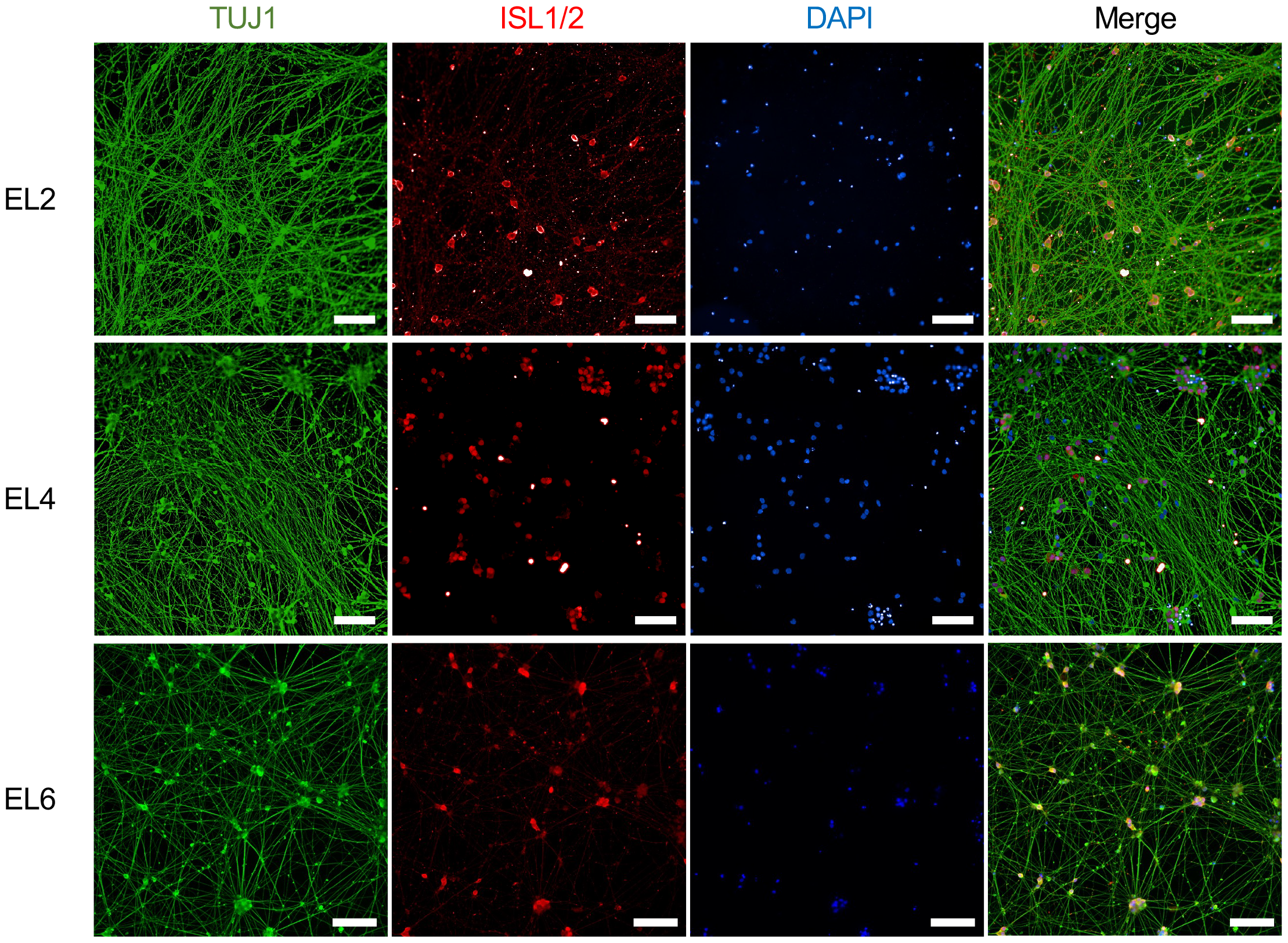
Forward programming of EL-specific iPSCs into cortical neurons. Representative images from EL iPSC lines brought through directed differentiation to forebrain cortical neurons. Maturation markers TUJ1 (green), ISLET1/2 (red), and DAPI (blue) are shown. Scale bars: 100 µM.

### Subject Consent and Global Distribution of Created Lines

All of the iPSC lines in this bank were created from subjects using a consent form under the Boston University Institutional Review Board (H32506). This consent form includes a comprehensive template that allows for the unrestricted sharing of deidentified created lines, including potential commercialization and sharing of lines with commercial entities. As a resource to investigators, this consent form has been included as <u>Supplemental Information</u>.

## DISCUSSION

Longevity is multifactorial and complex with a variety of environmental, lifestyle, and genetic determinants^45,46^. EL offspring demonstrate higher resistance to aging-associated diseases^5,7,47–50^ and they map younger in aging rate estimators than age-matched controls^51,52^. This last point is demonstrated in our cohort and in our application of a series of biological aging clocks. Biological aging clocks have emerged as a prominent tool for measuring and understanding the aging process at a molecular level^27,30,53,54^. Driven by advances in high-throughput sequencing, computational biology, and machine learning algorithms, researchers are continuously refining existing clocks, developing novel clock architectures, and expanding potential applications. In our study, we observed that many of the centenarians in our cohort demonstrated either significant differences in the comparison of biological to chronological age or slower rates of aging, even though studies have demonstrated that our rate of aging increases over a lifespan^29,55^. Moreover, genetic variants have been identified which are associated with longevity^8,46,56^ and resistance to aging-related diseases such as AD^57,58^. However, mechanistic insight into how these elements promote longevity remains speculative, highlighting the need for scalable human *in vitro* models that can be used to understand the gene and regulatory networks that promote resistance to or resilience against aging-related disease and inform the development of therapeutics which may slow or reverse aging.

As a step toward filing this gap in understanding, we have leveraged access to centenarians and their offspring in the New England Centenarian Study (NECS)^46^, the Longevity Consortium Centenarian Project (LCCP), and Integrative Longevity Omics (ILO) studies to build an EL-specific library of biomaterial from these exceptionally long-lived subjects. Using PBMC samples from these individuals, we demonstrated the ability to efficiently generate high-quality, fully pluripotent iPSCs regardless of subject age. This unique resource provides an unlimited amount of longevity-specific biomaterial (e.g., genomic DNA) and enables the generation of a multitude of cell and tissue types of aging-related interest to fuel longevity research and aid in the development and validation of novel geroprotective agents in a human model system. The ability to generate and assay multiple cell types is valuable, as tissue and organ systems have been shown to age at different rates^22,59^. Complementing the molecular profiling performed on the PBMCs of the subjects in our bank, we have curated associated demographics and cognitive and physical function characterizations of the subjects to highlight the relative health and functionality of the participants and their associated iPSC lines. These data should allow for more informed choices of which iPSC lines are best suited for particular research questions or screening applications in aging-related diseases such as neurodegeneration, where lines derived from those with resiliency or resistance to cognitive decline could serve as valuable models.

Interestingly, during the characterization process, although they were otherwise karyotypically normal, it was noted that a subset of the male centenarian iPSC lines displayed complete loss of the Y chromosome. This mosaic loss of Y (mLOY) has been previously identified in the PBMCs of males over 70 and may be a biomarker for aging and susceptibility to and prevalence for aging-related diseases such as cancer and cardiovascular disease^60–62^. Moreover, and conversely to that seen in centenarians, mLOY is also associated with a significant increase in all-cause mortality^61^. The inclusion of 3 male centenarians displaying mLOY in our bank allows for interesting research opportunities into the modeling and understanding of this phenomenon. Intriguing possibilities include the use of these particular centenarian lines to identify compensatory mechanisms against the deleterious impact of mLOY or the possibility that mLOY in these individuals is a beneficial adaptation for longevity. Additionally, we demonstrated that iPSC lines in this bank, including those with mLOY, are capable of competent differentiation to produce forebrain cortical neurons. As cortical neurons represent cell types of the central nervous system majorly impacted by the aging process and are intricately involved in aging-related decline, the ability to generate unlimited numbers of this target cell population from subjects who are potentially resistant to disease should prove to have great utility. Moreover, the inherent flexibility of an iPSC-based system also allows for the creation of a multitude of aging-impacted tissue types which can also be used in this capacity.

The use of iPSC-based platforms to study aging has received skepticism due to the loss of epigenetic information in resulting cells which is a byproduct of the reprogramming process^63^. Interestingly, this same process has gained interest in the context of rejuvenation, where the transient expression of the Yamanaka factors may return a cell to a more youthful state without the loss of cell identity, deemed ‘partial reprogramming’^64–66^. The epigenetic landscape and its associated changes across a lifetime have been identified as a hallmark of aging^45,67,68^. However, the genetics of an individual, which strongly impact longevity particularly at extreme ages^5,46,47^, are faithfully captured in iPSCs, enabling the potential discoveries that may arise from this level of information. Additionally, the reset iPSC epigenetic landscape presents a unique opportunity to simulate ‘aging in a dish’ by performing directed differentiations into distinct cell types, thereby reinitiating methylation changes and reinstalling a defined epigenetic landscape.

iPSC-based systems have revolutionized the modeling of genetic disorders and shown the ability to model diseases that manifest late in life^18–20,69–71^. These platforms allow for a variety of molecular studies to be performed, including those that employ novel gene editing tools to perturb specific genes and pathways associated with longevity. Here, we provide a unique resource of longevity-specific biomaterial that can be leveraged to build human *in vitro* models of aging-related disease and screen potential countermeasures. Models of *human* aging, longevity, and resilience to disease that allow for the functional testing of potential interventions are virtually non-existent. This resource directly addresses this limitation, while also allowing for cross-validation of the functional results, identified pathways, and observed signatures across other model systems and laboratories, a major point of concern in the rapidly emerging field of geroscience.

## METHODS

### Identification of subjects and curation of clinical history

Centenarians were identified from voter registries, mailings to adult living communities and long-term care facilities, news articles, and direct participation inquiries to the NECS, LCCP, or ILO studies. Offspring of living or deceased centenarians were also invited to participate. Spouses of the enrolled offspring were invited to participate as a referent group without familial longevity. Centenarian, offspring, and spouse participants complete self-administered questionnaires to collect sociodemographic, medical history, and physical function data. Participants also complete cognitive screeners and a neuropsychological and physical assessment by video conference or telephone. An informant reported on the presence of cognitive and psychiatric problems in the participant’s daily life. Comprehensive phenotypic data are available from the ELITE portal (https://eliteportal.synapse.org/). A convenience sample of NECS, LCCP, and ILO participants was selected for PBMC collection.

Functional independence for centenarians was defined as a score of 80-100 on the Barthel Activities of Daily Living Index (ADLs)^72^, a measure of independence in performing basic self-care. For the offspring and spouses, functional independence in performing independent living skills such as using a telephone and managing finances, known as Instrumental Activities of Daily Living (IADLs), was defined a score of 14 out of a possible score of 14 on the OARS Multidimensional Functional Assessment Questionnaire^73^. Cognitive status (i.e., cognitively healthy versus cognitively impaired) was determined by clinical consensus review of neuropsychological assessment scores or, if not available, from cognitive screeners (i.e., the Blessed Information Memory Concentration Test^74^ or the Telephone Interview for Cognitive Status^75^).

### Collection and isolation of PBMCs

Peripheral blood samples were procured from subjects with EL in the NECS^46^, the LCCP (https://www.longevityconsortium.org/), and the ILO Study (https://longevityomics.org/), including centenarians and their offspring. Additionally, age-matched spouse controls with no history of EL were collected as controls. In most cases, samples were immediately processed and frozen following isolation of PBMCs via Ficoll gradient. Peripheral blood was drawn and collected into BD Vacutainer™ Glass Mononuclear Cell Preparation (CPT) Tubes (BD 362761). These samples were then centrifuged at 1800 rpm for 30 minutes at room temperature (RT) to separate the blood plasma, PBMC buffy coat, and packed red blood cells. The buffy coat was extracted and washed with phosphate buffer saline (PBS) and centrifuged at 300 x G for 10 minutes at RT. The cell pellet was resuspended in PBS, counted, and cells were pelleted by centrifugation at 300 x G for 10 minutes at RT. Cells were resuspended at 8.0 x 10^6^ cells/mL in resuspension buffer and further diluted to 4.0 x 10^6^ cells/mL in freezing medium according to the 10X Genomics protocol (CG00039 Rev D). Cells were then transferred into cryovials and frozen at -80C before being transferred to long term -150°C storage.

### Multiparameter flow cytometry characterization of PBMCs

Cytometry characterization was performed according to OMIP-069 protocol 1^76^ detailing panel design, validation, and sample staining and acquisition. Frozen PBMCs were thawed rapidly and pelleted via centrifugation. Cells were stained with Live/Dead Fixable Blue dye (ThermoFisher L34961), washed, blocked with Human FcBlock (BioLegend 422301) and stained with a panel of 40 fluorescent reagents (**Supplementary Table 1**) supplemented with Monocyte Blocker (BioLegend 426102) and Brilliant Buffer Plus (BD Biosciences 566385). This cell mixture was allowed to stain for 30 min on ice and then washed. Ultracomp beads (ThermoFisher 011-2222-42) and control PBMCs were used to include single-stain controls. Cells and beads were then analyzed on a 5-laser Aurora spectral flow cytometer (Cytek Biosciences). At least 500,000 cells were recorded for each PBMC sample. Data were processed in SpectoFlo 2.2 (Cytek Biosciences) to generate unmixed features. OMIQ cloud platform was used to perform data cleanup and computational analysis. Live single CD45+ single cell datapoints were projected into two-dimensional space using PCA-informed opt-SNE dimensionality reduction algorithm^77^ and FlowSOM^78^ clustering of the same datapoints was overlaid to visualize biological population representation. FlowSOM metaclusters were annotated based on established phenotypic characteristics of the populations.

### Targeted methylation arrays and estimation of biological age

Isolation and purification of DNA: PBMCs were thawed rapidly, and DNA was extracted using the Qiagen DNeasy Blood and Tissue Kit (69506). Samples were submitted according to CD Genomics standards for Illumina Infinium MethylationEPIC v2.0 array.

### Methylation data quality control and pre-processing

Quality control metrics for all Infinium MethylationEPIC v2.0 samples were generated through the sesameQC_calcstats function in the SeSAMe R package (v1.20.0) on Bioconductor (v3.18)^79^. as detailed in their vignette. Beta values for each sample were obtained from idat files run through the openSesame pipeline from the same package as above, utilizing their recommended preprocessing for human EPICv2 samples. Preprocessing included quality masking for probes of poor design, dye bias correction, detection p-value masking, and background subtraction. Additionally, the EPICv2 array contains some duplicate CpGs across the 900k probes in its microarray, as well as additional suffixes added to probe IDs that reflect design information. To align these identifiers with those from models developed on earlier platforms and facilitate compatibility, the SeSAMe options collapseToPfx = TRUE, and collapseMethod = “mean” were utilized within the beta calling function, in order to remove suffixes, and average the betas from duplicate probes into a single value.

### Generating methylation clock estimates

Estimates from several high-profile age prediction models were performed with the methylation values generated from our samples. The models used were the original Horvath clock^27^, the Hannum clock^28^, a centenarian clock (ENCen40+)^31^, PhenoAge^30^, and DunedinPACE^29^. The Illumina EPICv2 platform comprises a new set of CpGs, which largely overlaps those found in EPICv1 and earlier platforms, but not entirely. As a result, each of these models had a small number of CpGs whose values were imputed to generate estimates as accurately as possible. For each model, imputation was followed as detailed in the corresponding manuscripts, if available. Otherwise, mean beta values were imputed from an external dataset belonging to the same Illumina platform on which a model was trained. For the Horvath 2013 model, approximately 13 of the 353 CpGs (3.68%) were missing from EPICv2, and missing values were obtained using mean DNAm values from a gold standard dataset detailed in the original supplement S2 (materials and methods) of the manuscript. Imputation with the centenarian clock (ENCen40+), as well as the PhenoAge clock, was done using mean DNAm values of the controls from a peripheral blood EPICv1 dataset GSE157252^80^ to supplement missing values for 39 of the 559 CpGs (6.98%) used by ENCen40+ and the 18 of 513 (3.51%) used by PhenoAge respectively. For the Hannum model, 7 CpGs of the 71 were missing (9.86%), and imputation was performed using the GSE40279 dataset (Illumina 450k) used in his original manuscript. Lastly, predictions for the DunedinPACE clock were generated using the Github repository (https://github.com/danbelsky/DunedinPACE) maintained by the authors. This repository has been independently updated to work with EPICv2 data, as the authors note that 29 of the 173 CpGs (16.76%) from the original model no longer appear in the new methylation array. The authors of this model utilize a similar approach of substitution of averages from Dunedin beta values for correction of missing data from their model, as well as for a panel of 19,827 probes for background normalization in the calculations.

### iPSC creation, expansion, and distribution

Previously isolated and frozen PBMCs were thawed rapidly and erythroblasts were expanded using erythroblast expansion medium (EM) for 9 days A comprehensive methodology concerning cell collection, expansion, and our approach to reprogramming can be found in Sommer et al., 2012. Following expansion, the cells were counted and 2.0 x 10^5^ cells were transduced using the Invitrogen™ CytoTune™-iPS 2.0 Sendai Reprogramming Kit (A16517). The next day, cells were collected and transferred onto matrigel-coated wells. Over the first 5 days post-transduction, the cells were slowly transferred to ReproTeSR (STEMCELL Technologies 05926) by adding the ReproTeSR on days 3 and 5 without aspirating the existing EM. From day 5 onward, the cells were cultured fully in ReproTesR. Once colonies adhered and were large enough, a cross-hatching method was used to divide each colony and passage onto a Matrigel-coated well to establish a clonal culture. These clones were expanded to at least 30 early passage vials and stored for sharing with the longevity community.

### Teratoma assay

3 female NU/NU immunocompromised mice (Jackson Labs strain#002019) per subject were injected subcutaneously with 1 x 10^6^ iPSCs (controlled for passage number) from each respective iPSC line suspended in high content Matrigel (Corning™ 354263) supplemented with 10uM Y27632 (Reprocell). Mice were monitored for teratoma formation for up to 20 weeks. Following teratoma development, the mass was resected, fixed in 4% PFA at room temperature overnight, and paraffin embedded and sectioned for hematoxylin and eosin staining.

### Forward programming of iPSCs to cortical neurons

Neurogenin 2 (NGN2) was expressed in iPSC lines using a tetracycline inducible promoter (Addgene 172115) integrated using Piggybac plasmid EFa1-Transposase (Addgene plasmid 172116) via lipofectamine (ThermoFisher STEM00015) as described previously^39^. Selection was performed after 48 hours using 2 μg/mL puromycin for 14 days. Stably integrated lines were dissociated with accutase (ThermoFisher A1110501) and plated at single-cell onto matrigel coated plates with induction media containing 10 μM Y-27632 and 2 μg/mL doxycycline (Sigma D9891) as previously described^40^. Following 3 days in induction medium containing doxycycline, cells were dissociated using accutase (ThermoFisher A1110501) and plated onto poly-L-ornithine (Sigma P4957), poly-D-lysine (Gibco A3890401), 10 μg/mL Fibronectin (Corning 356008) and 10 μg/mL Laminin (Gibco 23017015) coated plates in cortical neuron culture medium (CM) containing B27 (Gibco 17504044), BDNF (R&D Technologies 248-BDB) and NT3 (R&D Technologies 267-N3)^40^. Cells were patterned for 14 days in this medium before being used for downstream assays.

### Immunofluorescence

Cells were washed with PBS and fixed using 4% paraformaldehyde (Electron Microscopy Sciences 15714S) for 20 minutes at room temperature. Cells were permeabilized and blocked using 5% Fetal bovine serum (ThermoFisher 16141079), 0.3% Triton X-100 (Sigma T8787), and 2% bovine serum albumin (Gibco 15260037) diluted in PBS for 1 hour at RT. Cells were stained with primary antibodies (**Supplementary Table 2**) diluted in block/perm buffer overnight at 4°C. Cells were then washed 3 times with block/perm buffer and stained with corresponding secondary antibodies for 30 minutes at RT. Cells were washed 3 times with block/perm buffer and stained with Dapi nuclear stain (ThermoFisher 62248) diluted in fix/perm buffer for 10 minutes at RT. Cells were washed 3 times with block/perm buffer and imaged.

## Supporting information

Supplementary File 1. Consent for peripheral blood sample and associated patient data

## ACKNOWLEDGMENTS

GJM, TP, PS, and SA are funded by NIH-NIA (UH3 AG064704 U19 AG073172). Stephen Gayle, Lance San Souci, and Christos Meimeteas assisted with sample collection.

