## Supplementary Figures for "A longevity-specific bank of induced pluripotent stem cells from centenarians and their offspring"

**Supplementary Figure 1. Multi-dimensional flow cytometric characterization of the PBMCs of individuals with exceptional longevity.**

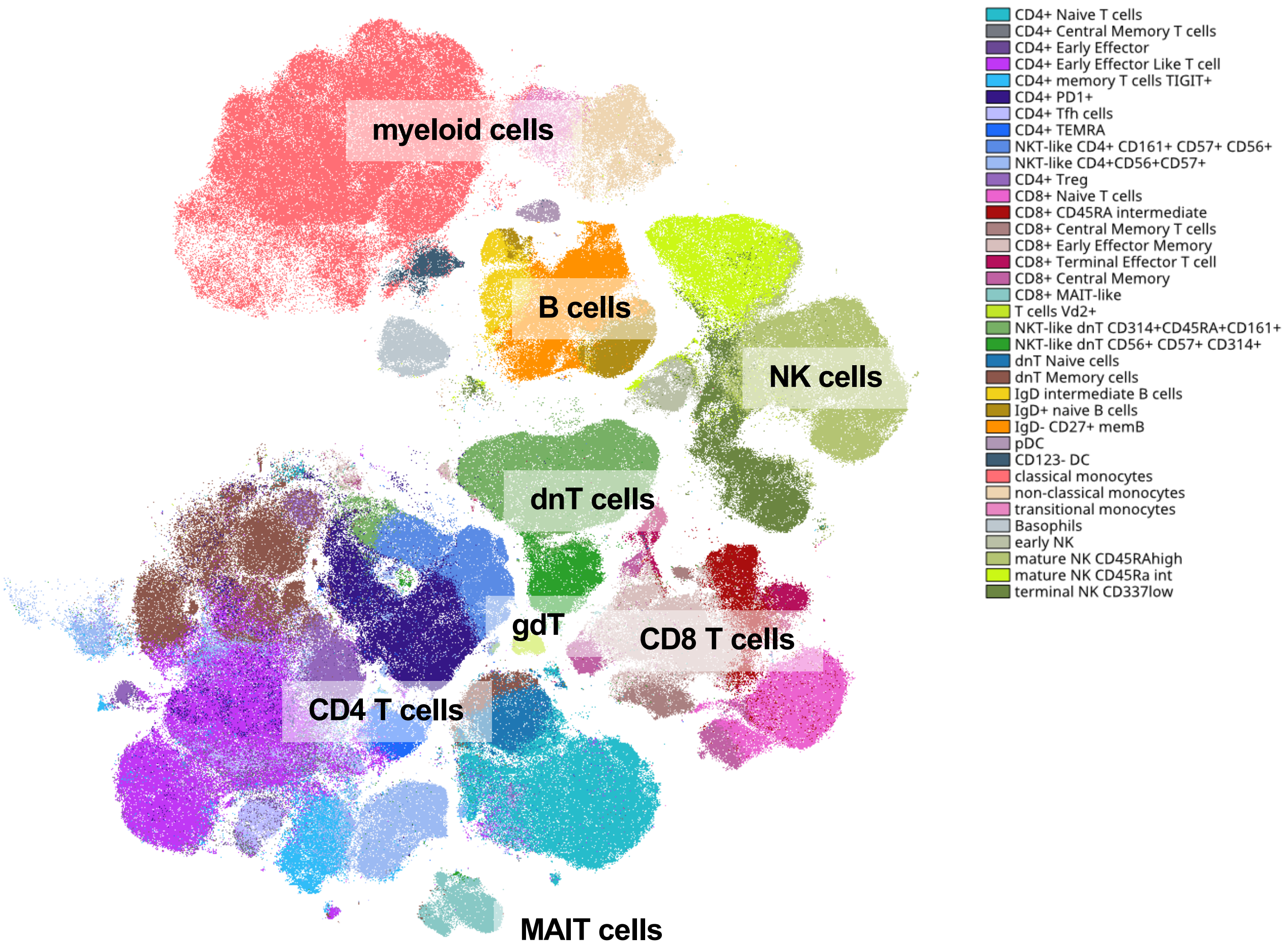


**Supplementary Figure 1. Multi-dimensional flow cytometric characterization of the PBMCs of individuals with exceptional longevity.** PCA-informed opt-SNE visualization of 40-color cytometry-characterized PBMCs with FlowSOM[82] clustering overlayed to annotate clusters based on established phenotype characteristics.

**Supplementary Figure 2. Phenotypic identification of PBMC sub-types within individuals with exceptional longevity.**

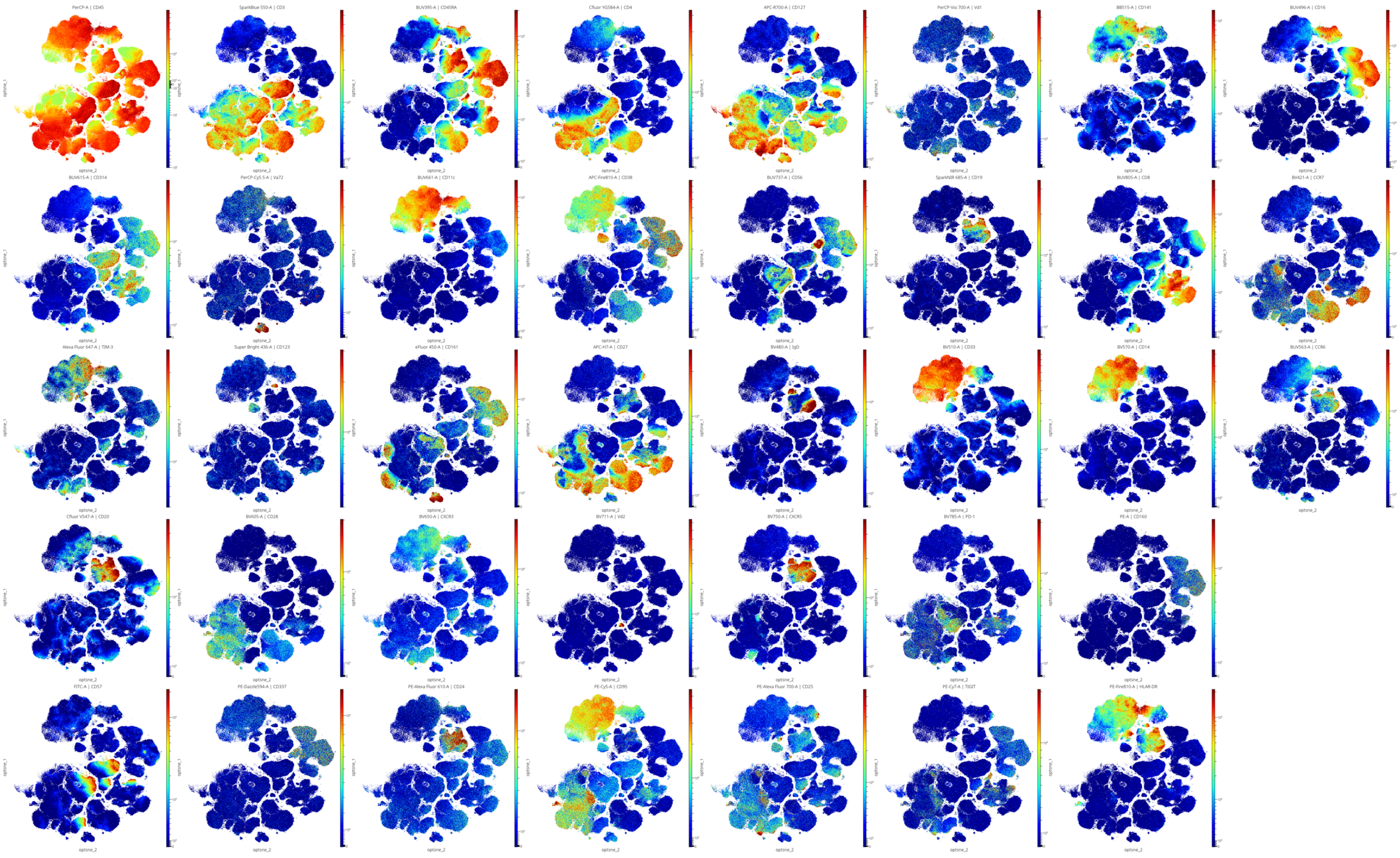


**Supplementary Figure 2. Phenotypic identification of PBMC sub-types within individuals with exceptional longevity.** Individual marker expression pattern overlayed on opt-SNE visualization of PBMCs based on established phenotypic characteristics.

**Supplementary Table 1. List of antibodies used for multiparameter flow cytometry characterization of PBMCs .**

| Ag | Fluor | Vendor | Cat No | amount per 1 test, ul |
|---|---|---|---|---|
| CD45RA | BUV395 | BD Biosciences | 740298 | 1.2 |
| CD16 | BUV496 | BD Biosciences | 612944 | 0.6 |
| CCR6 | BUV563 | BD Biosciences | 749362 | 3 |
| CD314 | BUV615 | BD Biosciences | 751232 | 5 |
| CD11c | BUV661 | BD Biosciences | 612967 | 1.2 |
| CD56 | BUV737 | BD Biosciences | 564447 | 1.2 |
| CD8 | BUV805 | BD Biosciences | 612889 | 1.2 |
| CCR7 | BV421 | BD Biosciences | 740052 | 5 |
| CD123 | SB436 | Thermo Fisher | 62-1239-42 | 2.5 |
| CD161 | eFluor 450 | Thermo Fisher | 48-1614-42 | 5 |
| IgD | BV480 | BD Biosciences | 566138 | 0.6 |
| CD33 | BV510 | Biolegend | 303422 | 2.5 |
| CD20 | PacOrange | Thermo Fisher | MHCD2030 | 5 |
| CD14 | BV570 | Biolegend | 301831 | 3 |
| CD28 | BV605 | Biolegend | 302968 | 2.5 |
| CXCR3 | BV650 | Biolegend | 353730 | 2.5 |
| Vd2 | BV711 | Biolegend | 331412 | 5 |
| CXCR5 | BV750 | BD Biosciences | 747111 | 1.2* |
| PD-1 | BV785 | Biolegend | 329930 | 5 |
| CD141 | BB515 | BD Biosciences | 565084 | 2.5 |
| CD57 | FITC | Biolegend | 359604 | 1.2 |
| CD3 | SparkBlue550 | Biolegend | 344852 | 2.5 |
| CD45 | PerCP | Biolegend | 304025 | 1.2 |
| Valpha7.2 | PerCPCy5.5 | Biolegend | 351709 | 2.5 |
| Vd1 | PCPVio700 | Miltenyi | 130-120-581 | 1 |
| CD160 | PE | Thermo Fisher | 12-1609-42 | 5* |
| CD4 | YG584 | Cytek Biosciences | R7-20042 | 1.2 |
| CD337 | PEDz594 | BD Biosciences | 325231 | 2.2 |
| CD24 | PEAF610 | Thermo Fisher | MHCD2422 | 2.5 |
| CD95 | PECy5 | Biolegend | 305610 | 0.6 |
| CD25 | PEAF700 | Thermo Fisher | MHCD2524 | 5 |
| TIGIT | PECy7 | Biolegend | 372714 | 1.2 |
| HLADR | PEFire810 | Biolegend | 900000155 | 5 |
| CD1d tetramer | APC | NIH tetramer core | | 0.02 |
| TIM-3 | AF647 | BD Biosciences | 565558 | 2.5 |
| CD19 | SparkNIR685 | Biolegend | 302269 | 1.2 |
| CD127 | APCR700 | BD Biosciences | 565185 | 6 |
| CD27 | APCH7 | BD Biosciences | 560222 | 2.5 |
| CD38 | APCFire810 | Biolegend | 303550 | 3.2 |

* reagents added to the cells before the rest of the reaction mix

**Supplementary Table 2. List of antibodies used for immunofluorescence.**

| Antibody | Vendor | Cat No | Dilution |
|---|---|---|---|
| Tra-1-81 | Thermo Scientific | MA1024 | 1:100 |
| Dapi | Invitrogen | D1306 | 1:1000 |
| Tuj1 | Millipore Sigma | T2200 | 1:200 |
| Islet1/2 | Abcam | ab86472 | 1:300 |
