## Supplementary File 1. Consent for peripheral blood sample and associated patient data for "A longevity-specific bank of induced pluripotent stem cells from centenarians and their offspring"

### RESEARCH CONSENT FORM

**Consent for peripheral blood sample and associated patient data**

GENERATION OF INDUCED PLURIPOTENT STEM CELLS (IPSC) FROM HUMAN SUBJECTS IN THE CREATION OF A REPOSITORY

#### Background

The purpose of this research is to create a repository or "bank" of a type of stem cell called "induced pluripotent stem cells" or iPSCs. Stem cells are special kinds of cells that can renew themselves. Under certain experimental conditions, iPSCs can be directed to grow into any type of tissue or organ. iPSCs can also live indefinitely in the laboratory. These stem cell lines will be used for medical research and/or to develop commercial research tools to create safer and more effective drugs.

You are being asked to provide a blood sample that will then be grown in the laboratory. You will also be asked for your permission for us to record some data from your medical record. Both the created cell lines as well as your medical record data may be released to other investigators or commercial entities in the future. Importantly, your name or other directidentifiers will never be released. The total duration of your involvement in this study is expected to be less than an hour.

The iPSCs that are created in this study will have many uses. For example, new drugs can be tested to see what effect they have on cells. Diseases can be "modelled" in a test tube so that we can have a better understanding of how they occur. New treatments for diseases can be tested using the cells. New tissues or organs could be created using iPSCs.

In addition to the immediate uses of the cells noted above, in the future iPSCs may be used in other ways. For example, research may include:

- Looking at the DNA sequence/genetic code in your cells
- Altering some of the DNA within these cells
- Testing in animals to model diseases and treatments

The iPSCs generated from your blood sample will never be used to clone (known as "reproductive cloning") or to otherwise create an entire human being. At the present time, research on your iPSCs is not likely to provide any information on your personal health. If your iPSCs do provide information about your personal health, we do not plan to provide you with this information. There may be a rare instance where we find information that we believe is urgently related to your health. In this very unlikely event, we may contact you to give you a choice about whether or not to learn the information.

This research consent form explains why this research study is being done, what is involved in participating in the research study, the possible risks and benefits of the research study,



*BMC and BU Medical Campus IRB*
*IRB NUMBER: H-32506*
*IRB APPROVAL DATE: 04/05/2023*

alternatives to participation, and your rights as a research subject. The decision to participate is yours. If you decide to participate, please sign and date at the end of the form. We will give you a copy so that you can refer to it while you are involved in this research study. We encourage you to take some time to think this over and to discuss it with other people and yourdoctor and to ask questions now and at any time in the future.

**Purpose**
The purpose of this study is to create a repository or 'bank' of induced pluripotent stem cells or iPSCs that can be used for future research. Scientists will use these iPSCs to study the biology of the disease or condition. It is hoped that in the future this repository will help lead to better treatments for genetic disease.

**What Happens In This Research Study?**
All or part of the research in this study will take place at Boston University Medical Center.

**Blood Sample:** You will be asked to provide a sample of blood which will be collected by a person trained in obtaining a blood sample. We will take approximately 16ml (4 teaspoons) of blood from your vein. The blood draw procedure will be performed once, unless the laboratory procedures fail, and in that case, an additional blood sample may be requested. The blood sample will then be taken to the laboratory to be made into iPSCs. The iPSCs will be frozen and saved for future research.

**Collection of medical information:** After the blood draw we will review your medical records and collect information about your past medical history. This information allows us to connect your clinical information with the created stem cell line, which is important for our studies.

The kind of information we will collect concerns things such as your disease or diagnosis, your family history, past physical examination results, and laboratory results. Importantly, sensitive information will also be collected including HIV status, sexually transmitted infections (STI) history, psychiatric history, and drug and alcohol history.

Additionally, we may contact you if we discover that the iPSCs made from your sample could be useful for research that is not covered by this consent form and that we want to get your permission to perform. This might include research on sperm and egg cells, reproduction, and infertility. It might include some research using new techniques or for new purposes that we simply cannot predict at this time. In the event the cell lines derived from your donation prove to offer a potential medical benefit, we may attempt to re-contact you to get additional health information if you agree by initialing below.

Please initial your choice:

_______________ I consent to being re-contacted in the future should the investigators wish to ask for additional health information.

_______________ I ***do not*** consent to being re-contacted in the future should the investigators wish to ask for additional health information.



***BMC and BU Medical Campus IRB***
***IRB NUMBER: H-32506***
***IRB APPROVAL DATE: 04/05/2023***

**Future Investigators Working with this Study or Conducting Other Research Studies:** We will keep your stem cell line and health data for future research. No information that can identify you will be shared with any other researchers or entities. We may give or sell this cell line and its derivatives and limited medical information about you to other researchers. These include other academic, non-profit and for-profit entities, including but not limited to hospitals, universities, cell/tissue storage banks, and businesses. Any sample which is shared with future investigators will ONLY be identified with a random code and it will not be possible to know whom this sample came from. Samples (blood cells, the stem cell line and its derivatives) obtained from you in this study may be used in the development of one or more diagnostic or therapeutic products which could be patented and licensed by those involved in the research or development of such products. There are no plans to provide financial compensation to you should this occur.

**What Are The Risks of This Study?**

**Blood Donation:** You may have some discomfort and bruising at the site of needle entry. There is a very small risk of fainting. Infection in the area of the needle insertion is rare.

**Emotional Risks Related to Potential Loss of Confidentiality:** Emotional and psychological risks are also possible with the donation of samples for iPSC creation. Patients uncomfortable with the thought of their cells being used for research purposes should not participate in the study. We may publish results of this research study in the medical literature.When we publish results, we do not use names or personally identifiable information. However,it is possible that you or family members could be recognized because of the rarity of your disease or based on your DNA sequence. It would be very difficult to identify any individual based on such published data, but it is a potential risk. Risk arises if your genetic information could be misused. For example, if research results suggested a serious problem with your health, it could be used to make it harder for you to get or keep a job or insurance. Although there is a small risk that your personal information will be released by mistake, there are laws in place that make it illegal for an employer or health insurance company to discriminate against an individual based on their genetic information.

**Group Risks:** Information on your ethnic and geographic background will be included with other medical information about you in the database as part of the stem cell bank. Research on the samples you provide may lead to results which are upsetting to you and others in your group, which you may disagree with, or which could be stigmatizing for your community.

**How Will My Confidentiality Be Protected?**

You have a right to privacy and we take reasonable measures to protect the confidentiality of your records and the information learned about you and your family members. Your name, birth date, and other personally-identifying information will be removed from your data and samples. They will be linked to your sample only by code number. The code key for the samples will be stored in a password-protected database under control of the Boston University School of Medicine investigators. Medical information, samples, and iPSC lines that are shared with others will be coded and will not include identifying information (name, address, telephone number, or personal identification number). Only the original investigators will be able to trace your samples and information to you. Information collected in this study may be reviewed by authorized individuals from the Food and Drug Administration (FDA), the



*BMC and BU Medical Campus IRB*
*IRB NUMBER: H-32506*
*IRB APPROVAL DATE: 04/05/2023*

National Institutes of Health (NIH), or other agencies for the purpose of making sure that proper systems, procedures, and regulations are being followed.

There is potential risk in genetic testing for uncovering and conveying unwanted information regarding parentage or specific risk of disease. Also, sometimes, knowledge of DNA test results can provoke anxiety and influence decisions regarding marriage and family planning: Both Massachusetts state laws and a new Federal law, called the Genetic Information Nondiscrimination Act (GINA), generally make it illegal for health insurance companies, group health plans, and most employers to discriminate against you based on your genetic information. These laws generally will protect you in the following ways:

1. Health insurance companies and group health plans may not request your genetic information that we get from this research.

2. Health insurance companies and group health plans may not use your genetic information when making decisions regarding your eligibility or premiums.

3. Employers with 6 or more employees may not use your genetic information that we get from this research when making a decision to hire, promote, or fire you or when setting the terms of your employment. All health insurance companies and group health plans must follow the GINA law by May 21, 2010. All employers with 15 or more employees must follow the GINA law as of November 21, 2009. Massachusetts law currently applies to all employers of 6 or more employees. Be aware that neither Massachusetts law nor the new Federal law protects you against genetic discrimination by companies that sell life insurance, disability insurance, or long•term care insurance. Thus, life insurance, disability insurance and long•term care insurance companies may legally ask whether you have had genetic testing and deny coverage for refusal to answer this question.

It is important to note that we will do everything we can to keep others from learning about your participation in this study. To further help us protect your privacy, we have obtained a Certificate of Confidentiality from the United States Department of Health and Human Services (DHHS). With this Certificate, we cannot be forced (for example by court order or subpoena) todisclose information that may identify you in any federal, state, local, civil, criminal, legislative, administrative, or other proceedings. The researchers will use the Certificate to resist any demands for information that would identify you, except to prevent serious harm to you or others, as explained below. You should understand that a Certificate of Confidentiality does not prevent you, or a member of your family, from voluntarily releasing information about yourself, or your involvement in this study. If an insurer or employer learns about your participation, and obtains your consent to receive research information, then we may not use the Certificate of Confidentiality to withhold this information. This means that you and your family must also actively protect your own privacy. Disclosure will be necessary, however,upon request of DHHS for the purpose of audit or evaluation, and is limited only to DHHS employees involved in the review. You should understand that we will in all cases, take the necessary action, including reporting to authorities, to prevent serious harm to yourself, children, or others. For example, in the case of child abuse or neglect.



*BMC and BU Medical Campus IRB*
*IRB NUMBER: H-32506*
*IRB APPROVAL DATE: 04/05/2023*

**Are There Benefits For Taking Part In This Study?**
Participation in this study will not benefit you or your family directly. However, your participation may help investigators to better understand certain diseases and conditions associated with your specific genetic variations, and to develop future safer and effective drugs and treatments for patients suffering from related conditions.

**Will I Receive Payment For Being In This Study?**
You will not be paid for taking part in this study. Your samples will be used for research, and they may also be used to make commercial products and treatments, meaning that they can be bought or sold in order to treat other people. The research done with your samples may help to develop new products in the future. You will not receive any financial compensation, should this occur.

**What If I Change My Mind And Do Not Want To Take Part In This Study?**
Donating your blood sample for this research project is completely voluntary. You have the right to not participate, and to refuse to provide your cells for this study. Your current or future medical care will not change in any way whether you agree or refuse to provide any cells for this research project. If you decide not to participate following the collection of your cells and/or the creation of your iPSC line, your cells and related medical information will bedestroyed. Importantly, in cases in which your iPSC line has been created and shared with other investigators, it may not be possible to destroy shared material.

**Your Rights**
By consenting to participate in this study you do not waive any of your legal rights. Giving consent means that you have heard or read the information about this study and that youagree to participate. You will be given a copy of this form to keep. If at any time you withdraw from this study you will not suffer any penalty or lose any benefits to which you are entitled. You may obtain further information about your rights as a research subject by calling the Office of the Institutional Review Board of Boston University Medical Center at 617-638-7207. The investigator or a member of the research team will try to answer all of your questions. If you have questions or concerns at any time, contact Dr. George Murphy at (617) 638-7541 during the day or Gustavo Mostoslavsky at (617) 459-3841 after hours.

**Protection of Subject Health Information**
You have certain rights related to your health information. These include the right to know who will get your health information and why they will get it. If you choose to be in this research study, we will get information about you as explained below.

HEALTH INFORMATION ABOUT YOU THAT MIGHT BE USED OR GIVEN OUT DURING THIS RESEARCH:

- Information from your hospital or office health records at BUMC/BMC or elsewhere. This information is reasonably related to the conduct and oversight of the research study. If health information is needed from your doctors or hospitals outside of BUMC/BMC, you will be asked to give permission for these records to be sent to the researcher.
- New health information from tests, procedures, visits, interviews, or forms filled out as part of this research study.



***BMC and BU Medical Campus IRB***
***IRB NUMBER: H-32506***
***IRB APPROVAL DATE: 04/05/2023***

- Anonymous information learned about your specific genetic polymorphisms of genes important to drug development or drug effects.

WHY HEALTH INFORMATION ABOUT YOU MIGHT BE USED OR GIVEN OUT TO OTHERS?
The reasons we might use or share your health information are:
- To do the research described here.
- To aid in developing better research tools for developing safer effective medicines and cell-based therapeutic treatments.
- To make sure we do the research according to certain standard set by ethics, law, and quality groups.

PEOPLE AND GROUPS THAT MAY USE OR GIVE OUT YOUR HEALTH INFORMATION
- PEOPLE OR GROUPS WITHIN BUMC/BMC
    - Researchers involved in this research study
    - The BU/BMC Institutional Review Board that oversees this research
- PEOPLE OR GROUPS OUTSIDE BUMC/BMC
    - People or groups that we hire to do certain work for us, such as data storage companies, or laboratories.
    - Agencies may include the U.S. Department of Health and Human Services, the Food and Drug Administration.
    - Administration, the National Institutes of Health.
    - Organizations that make sure hospital standards are met.
    - Other researchers that are part of this research study.
    - Other researchers that are not part of this research study.
    - Commercial entities using the created cell lines for profit
    - A group that oversees the research information and safety of this study.
    - Government agencies in other countries.

Some people or groups who get your health information might not have to follow the same privacy rules that we follow. We share your health information only when we must. We ask anyone who gets it from us to sign a confidentiality agreement to protect your privacy.
In most cases any health data that is being given out to others is identified by a unique study number and not with your name or other personal identifiers. So, although in some cases it is possible to link your name to the study data, this is not usually done.

**Are there other things you should know about?**
At the completion of this study, we plan to store any remaining sample for possible future use. The remaining samples and any cell lines created from the samples may be stored indefinitely by various researchers and entities and may be used for future studies, including genetic manipulation and other uses which cannot be predicted at this time. It is possible that derived cells or cell products may be placed into animals to study them as therapies.

**QUESTIONS:** If you have any questions, please do not hesitate to ask, and they will be answered at this time. If you think of any additional questions later about the study, contact the PI Dr. Murphy.



*BMC and BU Medical Campus IRB*
*IRB NUMBER: H-32506*
*IRB APPROVAL DATE: 04/05/2023*

**CONSENT:** I have read the information in this consent form describing this study (or have had it read to me). All my questions regarding the study and my participation in it have been answered to my satisfaction. I freely give my consent to participate in this study until I decide otherwise.

________________________________________________

**Subject (Signature and Printed Name) Date**

________________________________________________

**Legally Authorized Representative (Signature and Printed Name) Date**

________________________________________________

**Witness (Signature and Printed Name) Date**

________________________________________________

**Person Obtaining Consent (Signature and Printed Name) Date**



***BMC and BU Medical Campus IRB***
***IRB NUMBER: H-32506***
***IRB APPROVAL DATE: 04/05/2023***
